# Using Expansion Microscopy to visualize and characterize the morphology of mitochondrial cristae

**DOI:** 10.1101/2020.04.28.066605

**Authors:** Tobias C. Kunz, Ralph Götz, Shiqiang Gao, Markus Sauer, Vera Kozjak-Pavlovic

## Abstract

Mitochondria are double membrane bound organelles indispensable for biological processes such as apoptosis, cell signalling, and the production of many important metabolites, which includes ATP that is generated during the process known as oxidative phosphorylation (OXPHOS). The inner membrane contains folds called cristae, which increase the membrane surface and thus the amount of membrane-bound proteins necessary for the OXPHOS. These folds have been of great interest not only because of their importance for energy conversion, but also because changes in morphology have been linked to a broad range of diseases from cancer, diabetes, neurodegenerative diseases, to ageing and infection. With a distance between opposing cristae membranes often below 100 nm, conventional fluorescence imaging cannot provide a resolution sufficient for resolving these structures. For this reason, various highly specialized super-resolution methods including *d*STORM, PALM, STED and SIM have been applied for cristae visualisation.

Expansion Microscopy (ExM) offers the possibility to perform super-resolution microscopy on conventional confocal microscopes by embedding the sample into a swellable hydrogel that is isotropically expanded by a factor of 4-4.5, improving the resolution to 60-70 nm on conventional confocal microscopes, which can be further increased to ∼ 30 nm laterally using SIM. Here, we demonstrate that the expression of the mitochondrial creatine kinase MtCK linked to marker protein GFP (MtCK-GFP), which localizes to the space between the outer and the inner mitochondrial membrane, can be used as a cristae marker. Applying ExM on mitochondria labelled with this construct enables visualization of morphological changes of cristae and localization studies of mitochondrial proteins relative to cristae without the need for specialized setups. For the first time we present the combination of specific mitochondrial intermembrane space labelling and ExM as a tool for studying internal structure of mitochondria.

## Introduction

Super-resolution imaging has revolutionized fluorescence imaging by its capability to bypass the resolution limit of optical microscopy as defined by Ernst Abbe^1^. The most common methods, stimulated emission depletion (STED) microscopy^2^, photoactivated localization microscopy (PALM)^3^ and (*direct)* stochastic optical reconstruction microscopy (*d)*STORM^4^, were applied to countless biological specimens and can provide new insights into cellular structures and tissue in 2D and 3D^5, 6^.

However, super-resolution techniques enabling a resolution < 100 nm require specialized setups and expert knowledge to avoid artefacts^7^. Expansion microscopy (ExM)^8^ in contrast avoids this need by physical expansion of the sample after embedding into a polyacrylamide gel. Various protocols have been established using either a digestion or a denaturation step with numerous staining and linking protocols^9, 10, 11, 12, 13^. With an expansion of ∼4-4.5 times, ExM empowers scientists to resolve structures with a lateral resolution of ∼60-70 nm on a confocal microscope and in combination with structured illumination microscopy (SIM)^14^ of even ∼30 nm approaching the resolution of other conventional super-resolution methods^15^.

Many important biochemical processes take place in mitochondria, from the determination of cell fate by apoptosis induction, the citric acid cycle and the production of metabolites to the energy conversion *via* cell respiration. The latter is taking place at the mitochondrial inner membrane, which increases its surface by folding into cristae. The inner mitochondrial membrane can therefore be divided into the regions of the cristae membrane, which projects into the matrix, and the inner boundary membrane, which is found opposite to the outer mitochondrial membrane. Two regions meet at the so-called cristae junction^16^. Changes in morphology of cristae have been associated with ageing, numerous diseases, such as cancer, diabetes, several neurodegenerative diseases or types of neuro- and myopathies, and infection^17, 18, 19 20^. Thus, the possibility to investigate cristae morphology and the localization of mitochondrial proteins is of broad interest. Up to now, most light microscopy approaches have been performed using STED^21, 22, 23^ or Airyscan microscopy^24^. Although very successful in cristae visualization, the limitation is the restricted availability of super-resolution microscopes in standard cell biology laboratories as tools for investigating the mitochondrial ultrastructure.

Here, we report that ExM offers the possibility to image mitochondrial cristae on a classical confocal microscope and to localize mitochondrial proteins with an estimated lateral resolution of ∼30 nm in combination with SIM. We used green fluorescent protein (GFP)-labelled mitochondrial intermembrane space, mitochondrial creatine kinase MtCK-GFP as a cristae marker, and antibodies against mitochondrial matrix and cristae associated proteins. As an example of the applicability of this technique, using the combined resolution power of ExM and SIM we demonstrate that the mitochondrial transcription factor TFAM associates with cristae, and we observe changes in cristae morphology after membrane potential dissipation by CCCP or knockdown of the member of the mitochondrial intermembrane space bridging complex (MIB), Sam50.

## Material & Methods

### Cell culture

Human HeLa229 cells (ATCC CCL-2.1tm) and Sam50 knockdown cells *sam50kd-2*^25^ were cultured in 10% (v/v) heat inactivated FBS (Sigma-Aldrich, St. Louis, USA) RPMI1640 + GlutaMAXtm medium (Gibco, Thermo Fisher Scientific, Massachusetts, USA). The cells were grown in a humidified atmosphere containing 5% (v/v) CO_2_ at 37 °C. For the induction of the shRNA-mediated knockdown of Sam50 cells were treated with 1 µg/ml doxycycline for 72 h prior seeding.

### Transfection

MtcK gene was amplified from HeLa cDNA and cloned into the pCDNA3 vector (Thermo Fisher Scientific, Massachusetts, USA) where previously the GFP sequence was introduced, enabling C-terminal fusion and tagging. HeLa cells were transfected using Viromer® RED (230155; Biozym, Oldendorf, Germany) according to manufacturer’s instructions.

### Antibody conjugation

Following buffer exchange to 100 mM NaHCO_3_ with 0.5 ml 7 kDa Spin Desalting Columns (Thermo Fisher, 89882), the anti-TFAM (TA332462, rabbit; Origene, Rockville, USA) antibody was incubated in 5 molar excess of NHS-Alexa Fluor 546 (A20002; Thermo Fisher Scientific, Massachusetts, USA) for 3 h at RT. After conjugation, the unreacted dye was filtered from the antibody using 0.5 ml 7 kDa Spin Desalting Columns and the buffer was exchanged to 0.02 % NaN_3_ dissolved in PBS. The degree of labeling (DOL) was determined by the absorption of the antibody-dye with a UV-vis spectrophotometer (Jasco V-650). The labeled antibody was stored at 4 °C.

### Immunostaining

24 h after transfection, the cells were washed with 1xPBS and fixed with 4% PFA for 30 min at RT. Afterwards the cells were washed with 1xPBS, permeabilized for 15 min in 0.2% Triton-X100 and then blocked for 1 h in 2% FCS. Upon blocking, the cells were incubated for 1 h in primary antibody in a humidified chamber. We used the following primary antibodies: α-PRX3 (TA322472, rabbit; Origene, Rockville, USA), α-Mitofilin (ab48139, rabbit; Abcam, Cambridge, UK) α-TFAM (TA332462, rabbit; Origene, Rockville, USA) and α-GFP (ab1218, mouse; ; Abcam, Cambridge, UK). All primary antibodies were used in a dilution of 1:100. After incubation with the primary antibody, the cells were incubated with the secondary antibody, Alexa-488 (dilution 1:200, goat anti-mouse Alexa 488, Thermo Fisher Scientific, Massachusetts, USA) and ATTO647N (dilution 1:200, goat anti-rabbit ATTO 647N, 610-156-121S, Rockland Immunochemicals, Pottstown, USA). For 3 color images, the cells were then incubated in anti TFAM-Alexa 546 antibody (TA332462 self-conjugated or sc-166965; Santa Cruz Biotechnology, Dallas, USA).

### Expansion Microscopy

Expansion Microscopy was performed as described previously^9, 26^. Stained samples were incubated for 10 min in 0.25% glutaraldehyde, washed and gelated with a monomer solution containing 8.625% sodium acrylate (408220, Sigma-Aldrich, St. Louis, USA), 2.5% acrylamide (A9926, Sigma-Aldrich, St. Louis, USA), 0.15% N,N’-methylenbisacrylamide (A9926, Sigma-Aldrich, St. Louis, USA), 2 M NaCl (S5886, Sigma-Aldrich, St. Louis, USA), 1x PBS, 0.2% ammonium persulfate (APS, A3678; Sigma-Aldrich, St. Louis, USA) and tetramethylethylenediamine (TEMED, T7024; Sigma-Aldrich, St. Louis, USA,). Note that TEMED and KPS were added just prior to gelation and gelation was performed for 1 h at RT in humidified gelation chambers. The gelated samples were digested in the appropriate proteinase-containing buffer (50 mM Tris pH 8.0, 1 mM EDTA (ED2P, Sigma-Aldrich, St. Louis, USA), 0.5% Triton X-100 (28314, Thermo Fisher Scientific, Massachusetts, USA) and 0.8 M guanidine HCl (50933, Sigma-Aldrich, St. Louis, USA)) for 30 min and consequently expanded in ddH_2_O. The expansion factor was determined by the gel size prior and after expansion. Expanded gels were stored at 4 °C in ddH_2_O until use and immobilized on PDL coated imaging chambers (734-2055, Thermo Fischer Scientific, Massachusetts, USA) for imaging.

### Microscopes

Imaging of the unexpanded and expanded specimen was performed on a confocal system (Zeiss LSM700 5 mW red laser (637 nm) and a 10 mW blue laser (488 nm)) and on a structured illumination microscope (Zeiss Elyra S.1 SIM). In both cases a water objective (C-Apochromat, 63x 1.2 NA, Zeiss, 441777-9970) was used and images were processed with Imaris 8.4.1 and FIJI 1.51n^27^.

*d*STORM imaging was carried out on a home-built setup using an inverted widefield microscope (Olympus IX-71) equipped with an oil immersion objective (Olympus APON 60xO TIRF, NA 1.49) and an excitation laser of the wavelength 639 nm (Genesis MX639-1000, Coherent). The excitation beam was separated from the emitted fluorescence *via* a dichroic mirror (ZT405/514/635rpc, Chroma) and the emission was additionally filtered by an emission filter (Brightline HC 679/41 (Semrock)) in front of the EMCCD-camera (iXon Ultra 897, Andor). Prior to imaging, a switching buffer containing 100 mM ß-mercaptoethylamin pH 7.4 was added and 15 000 Frames at 50 Hz were recorded using laser densities of ∼ 7 kW/µm^2^. The super resolved images were reconstructed using the software rapidSTORM 3.3^28^.

## Results

Our first goal was to find a marker protein localizing to mitochondrial cristae, which can be expressed and correctly targeted when fused to a fluorescent protein without affecting mitochondrial function or morphology. After testing several candidates, we chose the MtCK, an enzyme providing a temporal and spatial energy buffer to maintain cellular energy homeostasis by creating phosphocreatine using mitochondrial ATP^29^. This protein localizes to mitochondrial intermembrane space, and after the transfection of HeLa229 cells and expression of its carboxy-terminus GFP-labelled version MtCK-GFP, we observed no effect on cell viability or mitochondrial length and distribution (not shown). The transfected Hela229 cells expressing MtCK-GFP were decorated with anti-GFP antibody prior to expansion, which we performed using the Chozinski protocol^9^ (Fig 1). Non-expanded and expanded cells were imaged by confocal fluorescence microscope (Fig 1a, c) and SIM (Fig 1b, d) and compared with *d*STORM images of unexpanded cells (Fig S1). Confocal imaging of expanded samples (Fig 1c) was comparable in resolution with the SIM of unexpanded cells (Fig 1b), whereas the combination of 4x ExM and SIM provided the best resolution of internal mitochondrial structure (Fig 1d).

**Figure 1:**
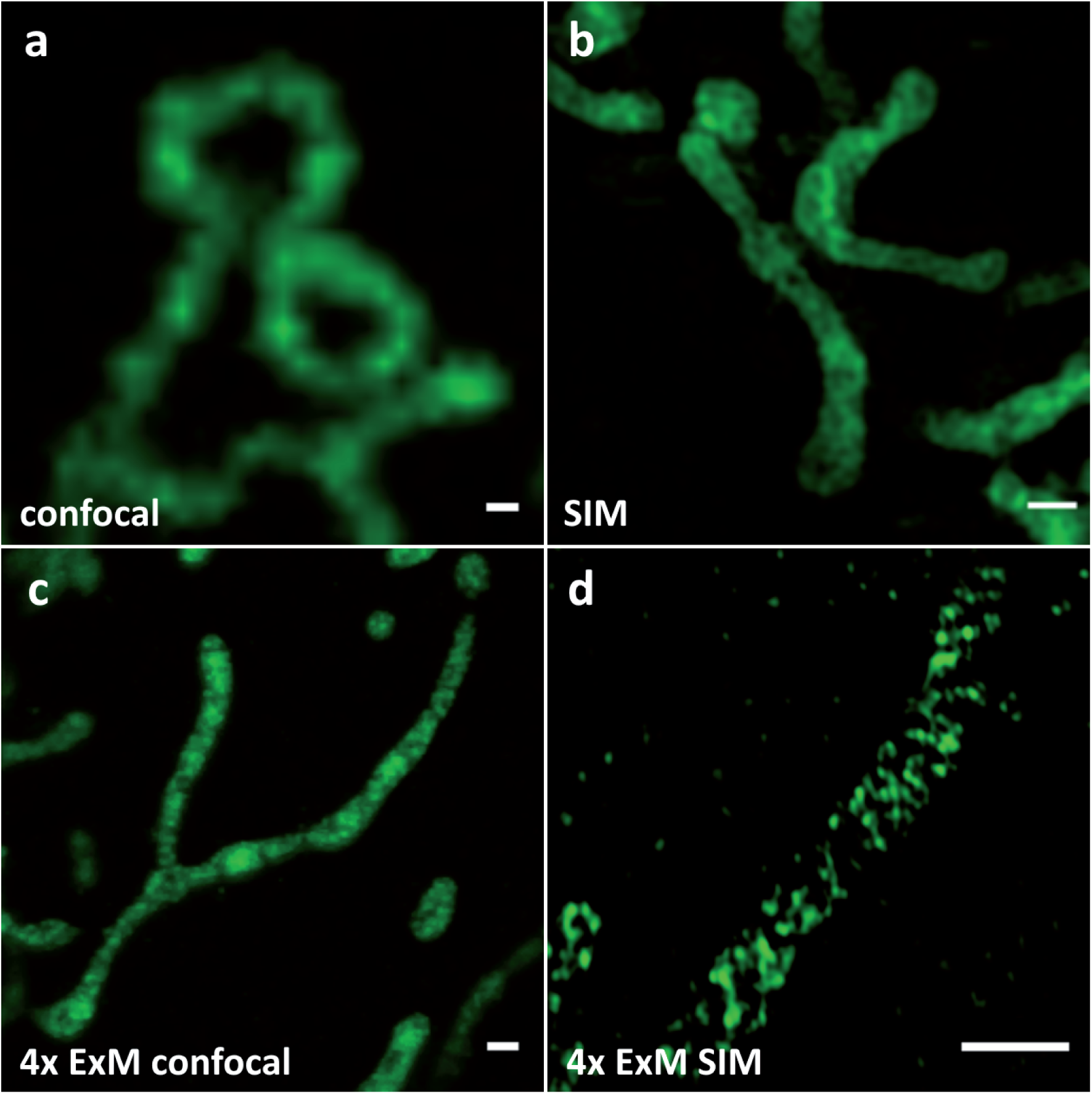
Comparison of (a) confocal, (b) SIM, (c) 4x ExM confocal (d) 4x ExM SIM images of Hela229 cells, transfected for 24 hours with MtCK-GFP (green) and immunolabeled for GFP to visualize mitochondrial cristae. Scale bars, (a,b) 0.5 µm and (c, d) 2 µm.

In addition to the GFP antibody labelling, MtCK-GFP-expressing cells were next decorated with antibodies against several mitochondrial proteins before expansion and analysed by confocal fluorescence microscope after 4x ExM (Fig 2). We chose peroxiredoxin 3 (PRX3), a mitochondrial thioredoxin-dependent hydroperoxidase present in mitochondrial matrix^30^, to assess the efficiency of MtCK-GFP to depict cristae with its uniform distribution through the intermembrane space (Fig 2a). This was indeed the case, since the signals for MtCK-GFP (green) and PRX3 (magenta) alternated and did not overlap, showing that 4x ExM enables the differentiation between the cristae and the matrix (Fig 2d). Next, we used the combined staining for GFP and Mic60/Mitofilin. The latter protein is one of the central components of the inner membrane mitochondrial cristae organizing system (MICOS), which together with the sorting and assembly machinery (SAM) in the outer mitochondrial membrane forms the mitochondrial intermembrane space bridging complex (MIB), a large complex necessary for the maintenance of mitochondrial cristae morphology, cristae junctions, inner membrane architecture, and formation of contact sites between two mitochondrial membranes^3131, 32, 33^. As we expected, we were able to localize Mic60/Mitofilin signals closer to the mitochondrial surface (Fig 2b) and overlapping with the signals of MtCK-GFP (Fig 2e). Finally, we used ExM to determine the localization of mitochondrial transcription factor A (TFAM) in relation to cristae. TFAM associates with mitochondrial DNA (mtDNA) nucleoids in the matrix, but a connection exists between cristae formation and mitochondrial nucleoid organisation^34^. Contrary to PRX3, TFAM is heterogeneously distributed in the matrix localized to punctate structures, which we observed overlapping with the MtCK-GFP signal (Fig 2c, f). This indicates that TFAM and mitochondrial nucleoids are found in the vicinity of cristae.

**Figure 2:**
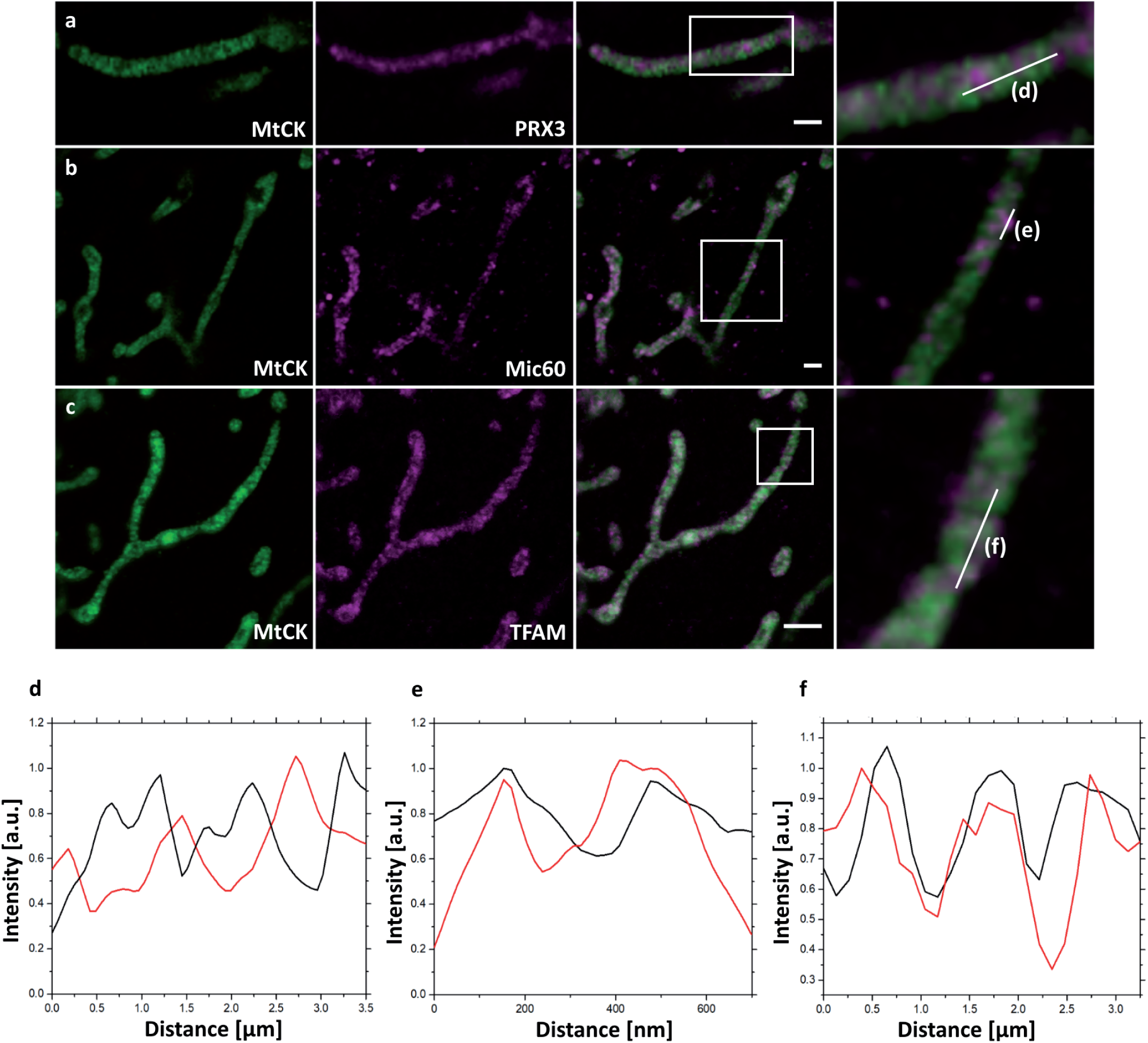
Confocal fluorescence images of 4x expanded HeLa229 cells transfected for 24 hours with MtCK-GFP (green) to visualize cristae, immunolabeled for GFP and (a) TFAM, (b) PRX3 or (c) Mic60/Mitofilin (shown in magenta). Plot profiling of the fluorescence of MtCK (black) relative to (d) PRX3, (e) Mic60/Mitofilin and (f) TFAM (red). 4x ExM can resolve if mitochondrial proteins localize at mitochondrial cristae as demonstrated for Mic60 and TFAM. Scale bars, 2 µm.

This finding was supported by SIM imaging of 4x expanded cells. The improved lateral resolution of ∼30 nm allowed us to visualize that most of the TFAM signal is found at the mitochondrial periphery, colocalizing with the MtCK-GFP signal, which indicates a possible association of TFAM and mtDNA nucleoids with cristae junction (Fig 3).

**Figure 3:**
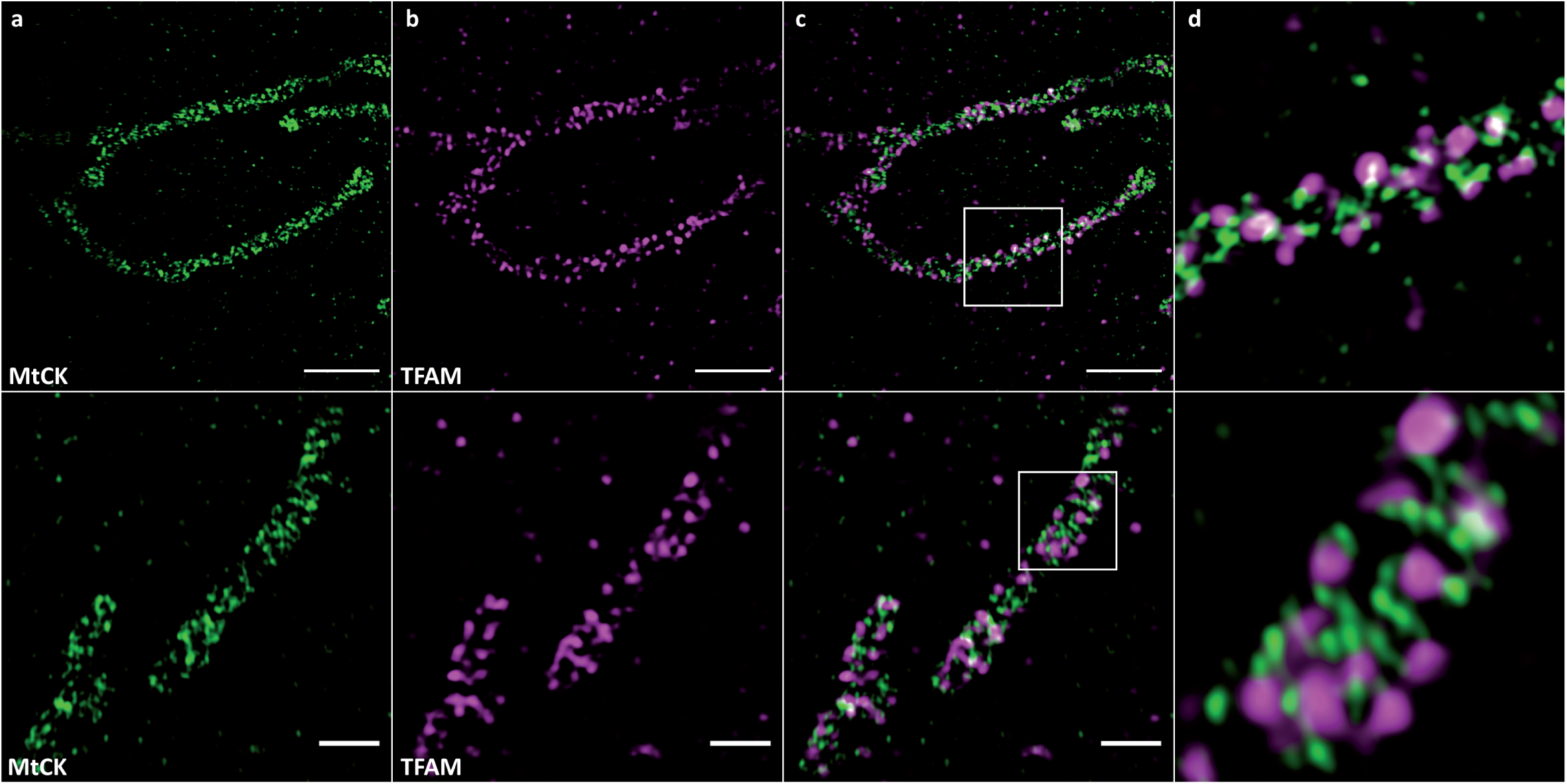
4x SIM-ExM to visualize the localization of TFAM relative to mitochondrial cristae. HeLa229 cells were transfected with MtCK-GFP for 24 hours, fixed, immunolabeled for GFP and TFAM and expanded. (a) MtCK-GFP (green), (b) TFAM (magenta), (c) merge and (d) zoom indicated in (c). SIM imaging further improves resolution and visualization of TFAM at cristae sites. Scale bars, 4 µm upper lane and 2 µm lower lane.

Next, we performed 3-colour confocal (Fig 4a-e) and SIM (Fig 4g-j) imaging of cristae, Mic60/Mitofilin and TFAM in expanded samples to better compare mitochondrial proteins and their localizations. The signals for all three proteins largely overlapped, but interestingly, the colocalization of Mic60/Mitofilin and TFAM was somewhat stronger than observed for Mic60/Mitofilin and MtCK-GFP (Fig 4f, k). This result confirms our previous observation that TFAM might localize in the vicinity of cristae junctions, which are marked by the presence of Mic60/Mitofilin.

**Figure 4:**
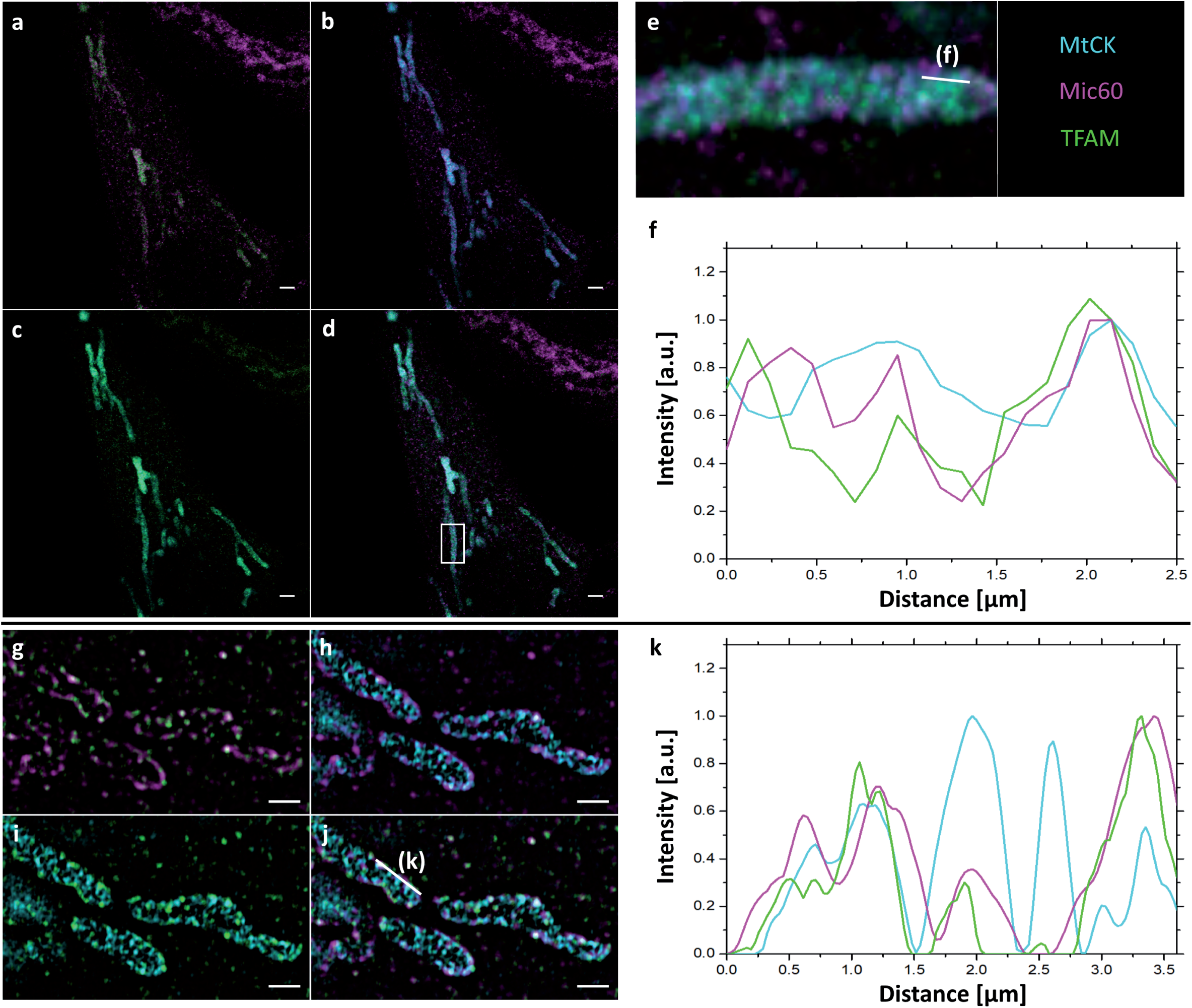
4x ExM of 3-color confocal (a-e) and SIM (g-j) images of the localization of mitochondrial proteins relative to mitochondrial cristae. Hela229 cells were transfected with MtCK-GFP for 24 hours, fixed, immunolabeled for GFP, Mitofilin and TFAM and expanded. (a, g) Mitofilin (magenta) and TFAM (green), (b, h) MtCK (cyan) and Mitofilin (magenta), (c, i) MtCK (cyan) and TFAM (green), (d, j) merge, (e) zoom from (d) with line indicating plot profiling shown in (f). (k) plot profiling line shown in (j). Scale bars, 5 µm for confocal images, 2 µm for SIM images.

To demonstrate the usefulness of 4x ExM for determining the morphology of mitochondrial cristae, we treated cells transfected with MtCK-GFP with 1 µM CCCP, a strong uncoupling agent that can abruptly depolarize the membrane potential ^35^. Cells incubated with CCCP exhibited a loss of cristae morphology in a time dependent manner, which was successfully visualized using MtCK-GFP and ExM (Fig 5a-d). Furthermore, we transfected HeLa cells where the knockdown of Sam50 could be induced by doxycycline (Dox)-mediated shRNA expression (*sam50kd-2*) with MtCK-GFP. Sam50 is a component of the SAM and MIB complex and, in addition to its function in the sorting of proteins with complicated topology, such as β-barrel proteins^36^, it is also important for the maintenance of cristae integrity. The enlargement of mitochondria and loss of cristae structure, that has already been visualized by transmission electron microscopy^31^, was also visible after ExM and analysis of the samples by confocal microscopy (Fig 5e, f).

**Figure 5:**
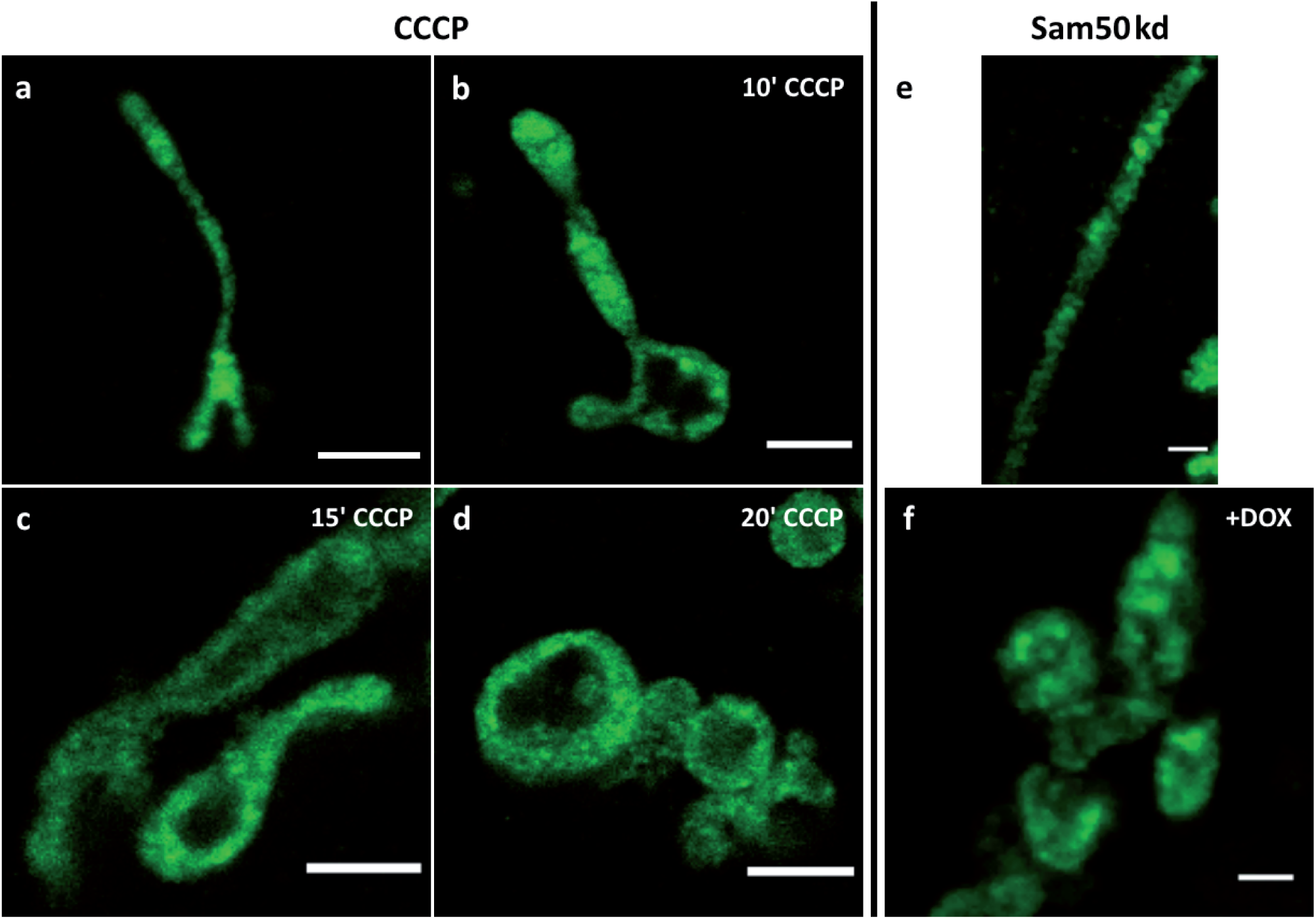
Confocal images of 4x expanded HeLa229 cells (a-d), transfected with MtCK-GFP (green), treated with 1 µM CCCP for 10 (b), 15 (c) or 20 (d) minutes. (e,f) Sam50 knockdown cells (*sam50kd-2*) were treated for 72 hours with 1 µg/ml doxycycline to induce Sam50 knockdown, transfected with MtCK-GFP (green) for 24 hours and analyzed by ExM on a confocal microscopy (f). The respective control sample without induced Sam50 knockdown is shown in (e). Scale bars, 2 µm.

## Discussion

In this study we could demonstrate the visualization of individual cristae on a conventional confocal microscope by applying ExM. This approximately fourfold expansion resulting in a resolution of ∼60 nm^8^ appears to be a promising tool for mitochondrial research, especially for cristae, folds of the inner membrane, with a distance often below 100 nm^37^. Imaging of cristae, however, has always been challenging due to the limited resolution in light microscopy. Hence, first successful results could be performed only with the advent of super resolution microscopy by applying SMLM^38^, STED^23^ or SIM^39^ approaches. While SMLM and STED have the limitation of highly specialized setups and intensive training, SIM has the drawback of an only to ∼ 100 nm limited resolution. Recently, studies showed great improvements in live-cell imaging using STED or Airyscan microscopy. These methods were used in combination with fluorescent dyes or inner membrane markers for analysis of the mitochondrial ultrastructure^21, 22^, measurement of the membrane potential of individual cristae^24^ or even cristae dynamics^40^. Although these live-cell imaging methods enabled following of cristae dynamics in real time, they also exhibited very fast damaging of mitochondrial vitality and are also not suitable for protein localization studies.

Contrary to those already established cristae marker COX8A^22^ or ATP5I^40^, we assume that we do not interfere with the naïve ATP production cycle using the mitochondrial creatine kinase MtCK as cristae marker. Other fluorescent dyes like MitoPB Yellow^21^ or MitoTracker for labeling cristae were unfortunately not compatible with ExM due to the lack of a primary amine, making MtCK a preferred possibility to label cristae.

Our approach by fixing and expanding the sample offers the possibility of immunolabeling and thus enables studies of protein localization relative to mitochondrial cristae with an estimated resolution of 60 nm. By additionally applying SIM the resolution is improved to ∼30 nm (Fig 3, 4g-j), approaching super-resolution methods like *d*STORM (Fig S1) and STED. Moreover, ExM also offers the possibility of super-resolved imaging of 3 or even 4 colours of the whole cell, a severe obstacle in super-resolution microscopy^32^.

In our study, we successfully demonstrated the close proximity to cristae of Mic60/Mitofilin, a protein localized to cristae junction and essential for cristae formation^32, 41^, and for the first time TFAM, a mitochondrial transcription factor located in the mitochondrial matrix and associated with mtDNA nucleoids. In comparison to the localization relative to cristae of TFAM and another mitochondrial matrix protein PRX3 a clear difference can be observed. While TFAM co-localizes with cristae, PRX3 shows an alternating signal (Fig 2). 3-colour imaging (Fig 4) and ExM in combination with SIM (Fig 3, 4g-j) indicated the possible localization of TFAM to cristae junction, which would explain the observed changes in mtDNA nucleoids upon Mic60/Mitofilin depletion^34^, and would also be in agreement with our unpublished data, which show a strong reduction in the quantities of mitochondrial genome maintenance exonuclease 1 (MGME1/C20orf72) upon Mic60/Mitofilin knockdown. MGME1 is an exo- /endonuclease involved in the maintenance of proper 7S DNA levels in mitochondria and mtDNA repair^42, 43^. It is therefore possible to imagine that the changes in mitochondrial morphology are sensed and responded to through the positioning of nucleoids at cristae junction and the possible functional connection of some of the mtDNA-associated proteins with the MICOS complex. However, more evidence needs to be obtained to support this theory, for which ExM of mitochondria might prove to be a useful tool.

Finally, we used ExM of mitochondria with MtCK-GFP labelled cristae to visualize the defects in cristae morphology after treatment with CCCP (Fig 5), which dissipates membrane potential and leads to the swelling of mitochondria and loss of cristae structure. With increasing length of incubation, we could observe successfully the loss of cristae integrity, which is in accordance with previous studies. Using our tool, we could also observe the loss of cristae after knockdown of Sam50 as reported before^31^ further demonstrating the usefulness of the MtCK-GFP as a marker for investigating cristae morphology. The combination of cristae labelling and ExM we present here is therefore a useful and relatively simple method for monitoring changes in cristae structure upon various stimuli, which could include up- or downregulation of proteins, chemical treatment or infection. Also, this tool can be used for determining submitochondrial localization of novel proteins in regard to cristae, enabling simultaneous immunodecoration for multiple markers and requiring only the usage of widely available fluorescent confocal microscopy.

## Acknowledgments

We thank Thomas Rudel and Georg Nagel for scientific input and for critically reading the manuscript.

## Data Availability

The raw data supporting the conclusions of this manuscript will be made available by the authors, without undue reservation, to any qualified researcher.

## Author Contributions

TK, RG, MS and VKP conceived the study, wrote the manuscript, and edited the manuscript. TK and RG performed the experiments and analyzed the data. SG performed the cloning of the construct. VKP and MS supervised the study.

