## Supplementary figures and images for "Using Expansion Microscopy to visualize and characterize the morphology of mitochondrial cristae"

### Supplemental Figure 1

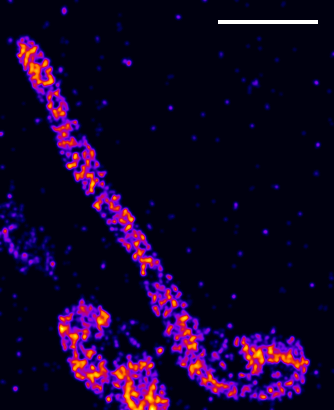
