## Supplemental Figure Legend for "Using Expansion Microscopy to visualize and characterize the morphology of mitochondrial cristae"

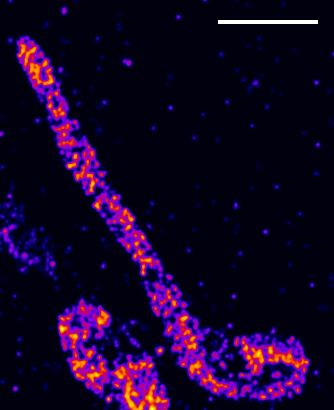

**Supplemental Figure S1:** dSTORM image of HeLa229 cells transfected with MtCK-GFP for 24 hours, fixed and immunolabeled for GFP. Scale bar, 2 µm.
